# VSIM: Visualization and simulation of variants in personal genomes with an application to premarital testing

**DOI:** 10.1101/529461

**Authors:** Azza Althagafi, Robert Hoehndorf

## Abstract

**Background:** Interpretation of personal genomics data, for example in genetic counseling, is challenging due to the complexity of the data and the amount of background knowledge required for its interpretation. This background knowledge is distributed across several databases. Further information about genomic features can also be predicted through machine learning methods. Making this information accessible more easily has the potential to improve interpretation of variants in personal genomes.

**Results:** We have developed VSIM, a web application for the interpretation and visualization of variants in personal genome sequences. VSIM identifies disease variants related to Mendelian, complex, and digenic disease as well as pharmacogenomic variants in personal genomes and visualizes them using a webserver. VSIM can further be used to simulate populations of children based on two parent genomes, and can be applied to support premarital genetic counseling. We make VSIM available as source code as well as through a container that can be installed easily in network environments in which genomic data is specially protected. VSIM and related documentation is freely available at https://github.com/bio-ontology-research-group/VSIM.

**Conclusions:** VSIM is a software that provides a web-based interface to variant interpretation in genetic counseling. VSIM can also be used for premarital genetic screening by simulating a population of children and analyze the disorder they might be carrying.

## Introduction

The contribution of genetics in human disease may range from almost 100% for monogenic, Mendelian disorders to a much smaller percentage for complex diseases including infectious disease [1]. Understanding how variation in individual’s genome relate to disease risk is important as it allows us to prevent and predict health effects in individuals, generate better diagnoses and prognoses of disease, and enable new approaches for treatment and development of the new drugs [2]. Predicting possible health effects from genome sequences is a significant emerging challenge [3] and important to support genetic counseling and preventing major health problems. The interpretation of whole-exome or whole-genome sequencing data linked to individuals is increasingly being used to identify causal mutations that may lead to an abnormal phenotype or a disease [3]. Visualization is one of the means to interpret genomic data and it plays a crucial role in data analysis [4].

While the interpretation of variants in individual genomes can be used to make personal health choices, it is also a challenge to determine the risk of particular phenotypes in children from two individuals. Consanguineous marriage is common in some regions of the world as a result of socio-cultural factors including religion and ethnicity [5, 6, 7, 8]. Recent studies show that the prevalence of consanguineous marriages is varying from 33–68% in different countries [5], and is estimated around 58% in Saudi Arabia [9], 50% in United Arab Emirates [10], 50% in Oman [11], and 68% in Egypt [12]. As a response to the health challenges arising from high rates of consanguinity, genetic testing and screening is introduced by governmental health authorities with voluntary or mandatory genetic testing before marriage to identify individuals who are carriers of autosomal recessive disorders, or individuals which have a genetic predisposition that may produce a disease in their children [5, 6, 7].

For example, the Saudi Premarital Screening and Genetic Counseling (PMSGC) program named the “Healthy Marriage Program” is part of a national project led by the Saudi Ministry of Health [13]. There are a comprehensive PMSGC program guidelines distributed to all workers in the program. Couples with marriage proposals are required to report to the nearest health-care clinic to apply for premarital certificates [14]. This test checks for specific diseases, in particular hemoglobinopathies such as sickle cell anemia and thalassemia, and infectious diseases such as infection with human immunodeficiency virus (HIV), hepatitis B virus (HBV), or hepatitis C virus (HCV) [15]. However, the implementation of premarital infectious disease screening that captures more diseases is an ambitious and extensive project [16, 13], and while infectious disease screening is important, it does not address the problem of heritable diseases arising from consanguinity.

There are several large public databases that contain information about the likely phenotypic effects of variants, including their penetrance and effect sizes [17]. Furthermore, several methods have been developed that can predict likely phenotypic effects of variants using a wide range of features including evolutionary conservation, protein structure and function, network connectivity, and likely association of a variant and a phenotype [17]. These methods already provide rich information that can be used to interpret individual genomes. However, while Mendelian diseases may be predicable in children based on the genome sequences of the parents using Mendel’s laws of inheritance, this is not true for more complex diseases including di- and oligogenic diseases; partially, their inheritance is affected by linkage disequilibrium that results in an non-uniform distribution of recombination centered around recombination hotspots.

We have developed VSIM as a tool for the visualization and interpretation of likely disease-causing variants in individual WGS or WES sequences. Given two genomic sequences (WGS or WES) as input, VSIM is further capable to simulatea cohort of child genomes, taking into account the recombination rates and probabilities. We use the information about members of this cohort of child genomes to determine the probabilities that children of the two individuals from which the original genomic sequences were derived will develop a certain disease or phenotype. VSIM can therefore not only be used to interpret and visually explore individual genome sequences, but also to perform premarital genetic testing. VSIM relies on information about Mendelian diseases from the ClinVar database[18], genetic associations for risk factors and complex diseases from Genome-Wide Association Studies (GWAS)[19], digenic disease variants from the DIgenic disease DAtabase (DIDA) [20], as well as pharmacogenomic variants from the Pharmacogenomics Knowledgebase (PharmGKB) [21]. Optionally, VSIM can also include the Mendelian Clinically Applicable Pathogenicity (M-CAP) Score [22]. The output of VSIM relies on Ideograms [23] that are easy to interpret and understand and from which additional information about the variant and its likely phenotypic effect can be accessed. VSIM is freely available as source code and as a Docker container at https://github.com/bio-ontology-research-group/VSIM.

## Result

VSIM is web-based simulation and visualization tool which aims to support genetic counseling and interpretation of genomic sequences. VSIM performs two main operations: first, VSIM is able to annotate and visualize personal genomes available in the VCF file format [24] in order to support visual exploration of variants and other genomic aberrations that may have an impact on health. Second, given two VCF files for two potential parents, VSIM can simulate a population of children, based on accurately accounting for recombination probabilities across the human genome, and then allows visual exploration of the simulation results. One of the main applications of the second feature of VSIM is genetic counseling and premarital genetic testing, but the simulation and annotation of genomes can also be used for evolutionary studies.

### Annotating and Visualizing Personal Genomic Data

VSIM accepts a VCF file as input, annotates the variants in the VCF file, and then visualizes the results on a chromosomal ideogram. Annotation of variants falls into five categories: known Mendelian disease variants (using the information from the ClinVar database [18]); disease-associated variants derived from GWAS studies (using the information from the GWAS Catalog [19]); variant combinations in digenic disease (using the information provided by the DIDA database [20]); pharmacogenomic variants (from the PharmGKB database [21]); and predicted pathogenic variants (using the M-CAP pathogenicity score [22]).

VSIM then generates chromosomal views based on chromosomal ideograms and shows the chromosomal positions at which functional variants have been found; this chromosome-focused visualization allows, for example, identifying haplotype blocks that are enriched for functional variants. Different categories of variants are shown in different colors, and it is possible to filter variants by their type (e.g., whether they are Mendelian disease variants, pharmacogenomic variants, etc.). Users are able to obtain additional information about variants when selecting a single variant, and can follow a hyper-link to a website with additional information and evidence about the type of variant. Figure 1 provides an example of the visual output produced by VSIM from a single VCF file.

**Figure 1:**
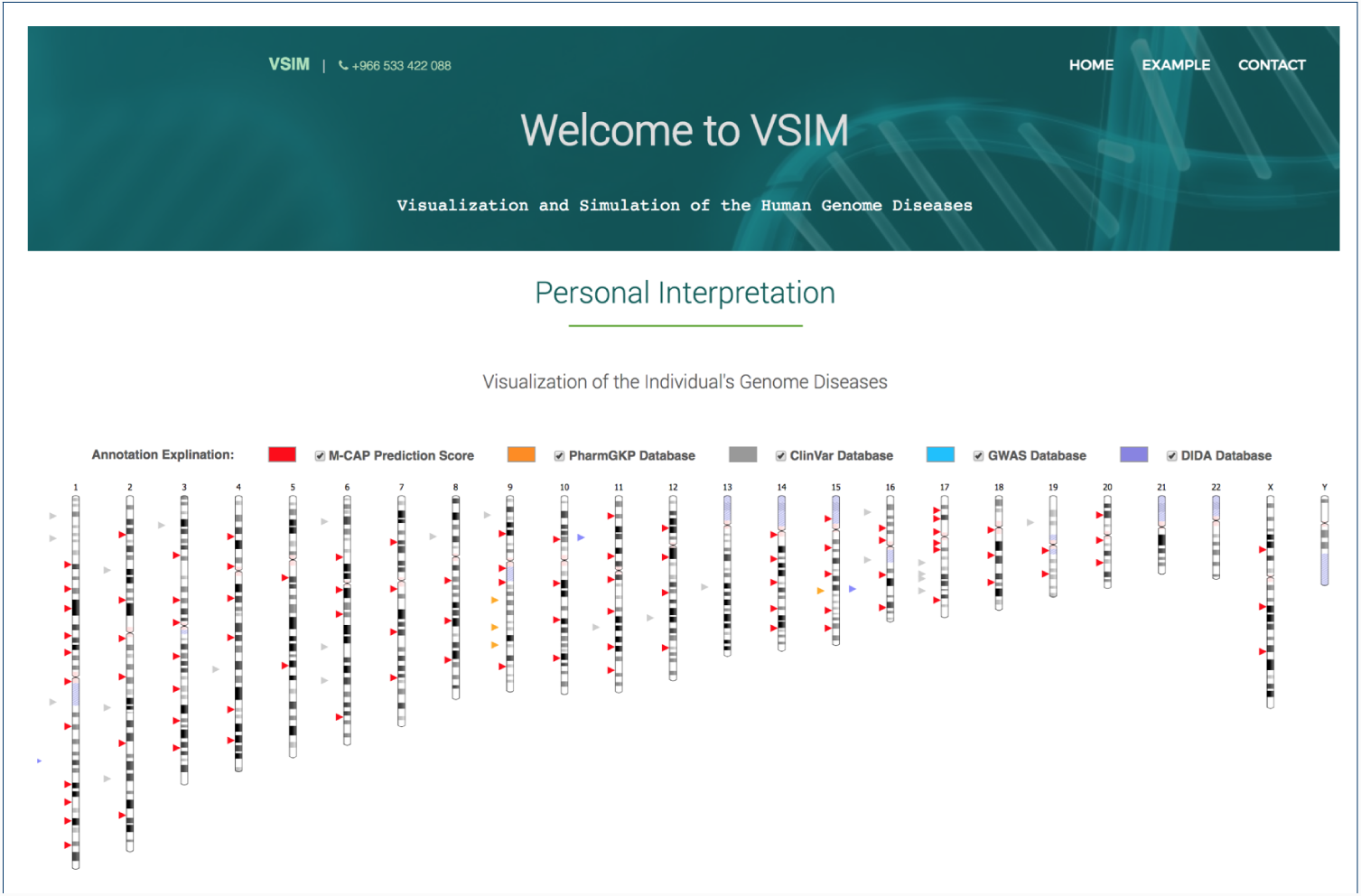
Visualization of individual genomes

### Simulating Child Cohorts and Application to Premarital Testing

VSIM is further capable of simulating cohorts of potential child genomes when given two VCF files as input, and using this simulated cohort to estimate the probability of encountering particular genetically based diseases in potential children (as well as the co-morbidities between the diseases). For this purpose, VSIM uses a map of genome-wide recombination rates for the human genome [25] which provides a global (i.e., not population-specific) estimate of recombination rates, distinguished by male and female genomes. The recombination rate is derived from 3.3 million crossovers from 104,246 meioses (57,919 female and 46,327 male meioses) [25].

Using the two input VCF files, the recombination rates and a parameter that determines the number of cross-overs per chromosome, VSIM simulates a population of potential children while considering the recombination probabilities; therefore, the population of children will account for, at least partially, linkage disequilibrium and the resulting correlation between risk-conveying or causative genomic positions. We annotate all genomes in the simulated cohort of children using the same annotation procedure and annotation sources used by VSIM, and we use the percentage of children within the population that carries a particular functional variant to estimate the likelihood that children develop a particular disease. While the likelihood could be estimated directly using Mendel’s laws from the two parent genomes in Mendelian diseases, our simulation approach will give more accurate probabilities in the case of complex, digenic, and oligogenic diseases, and further allows the estimation of co-morbidities in the child population. Figure 2 provides an example of the simulation result and its visualization.

**Figure 2:**
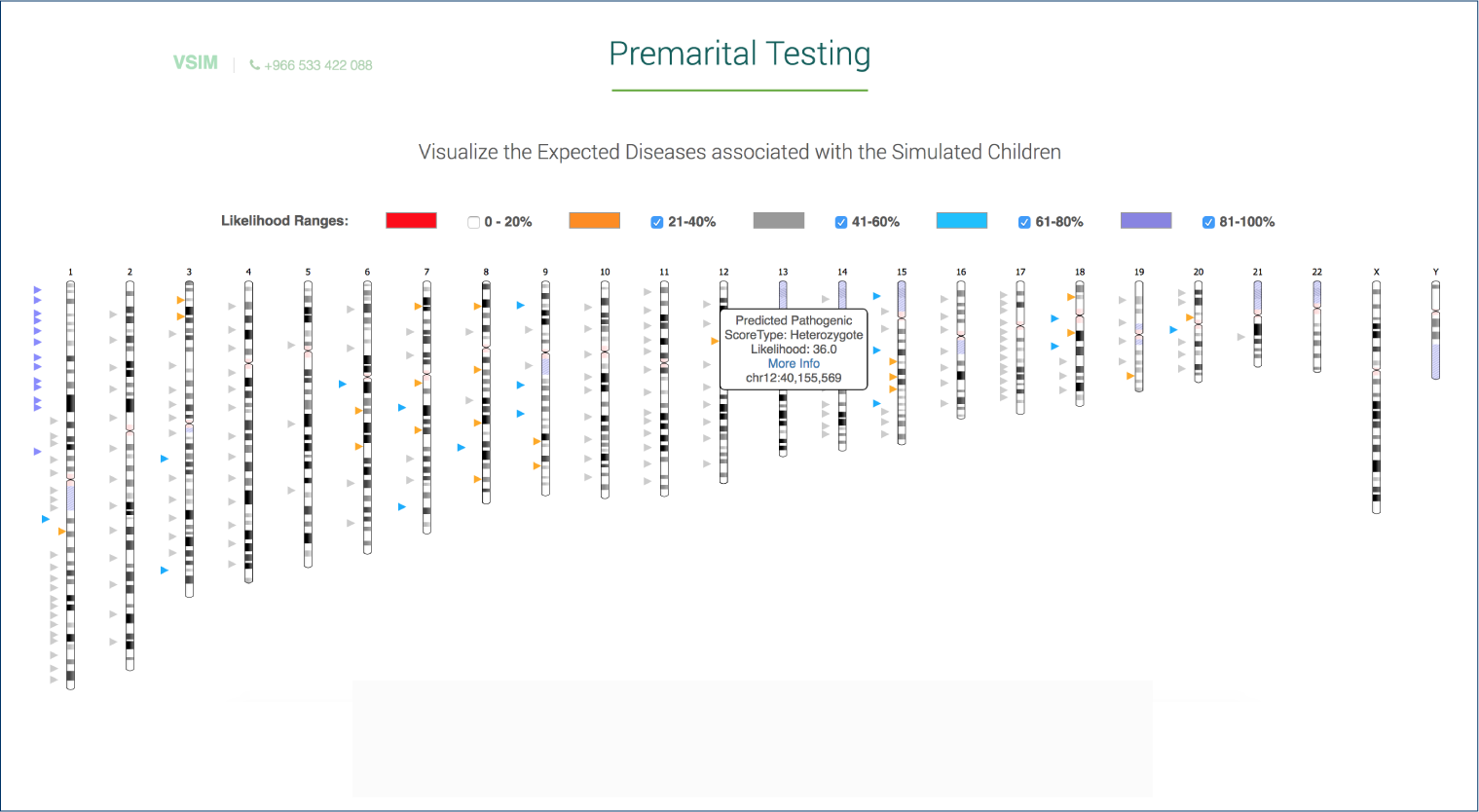
Example of Simulation result

### Evaluation

The time it takes to analyze (i.e., annotate and visualize) a single whole genome depends on the size of the VCF file. For a VCF file with 3 million SNPs, VSIM needs approximately 10 minutes to generate the final output using an Intel i7 processor at 2.5GHz with 16GB of memory. VSIM annotates a single variant on average in 1.4 × 10^−4^ seconds.

When applying VSIM for determining the likelihood of children having or carrying a particular disease, the simulation time not only depends on the size of the VCF file but also depends on the number of simulated children. Figure 3 shows the performance benchmarks for a different number of simulated children. The time linearly increases with the number of simulations to perform, and the generation of simulated genomes can easily be parallelized.

**Figure 3:**
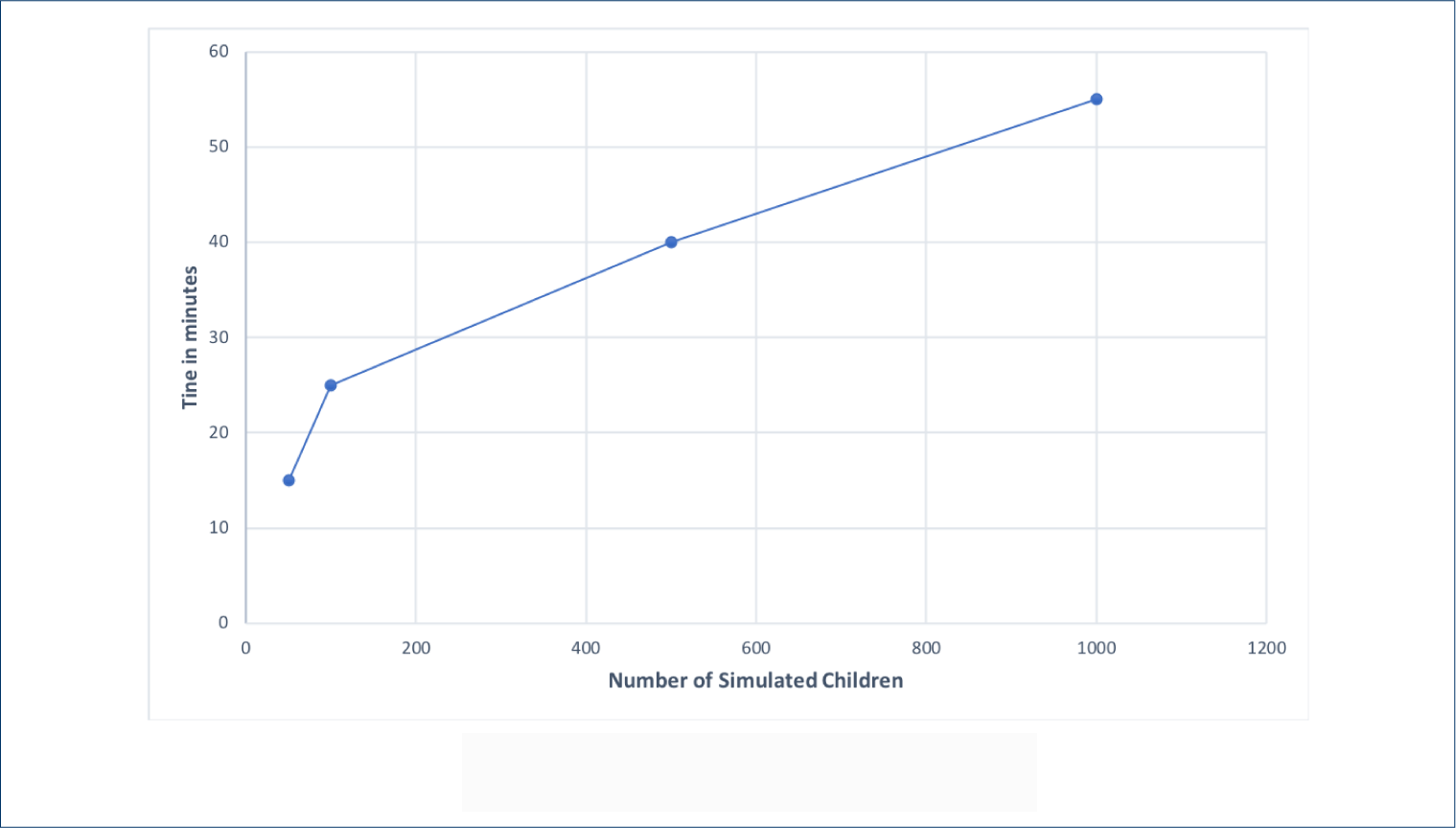
Simulation time

In addition to time, we also estimate the number of generations our simulator needs to produce linkage disequilibrium as observed in a real population. We start with a randomly generated population of individuals and randomly pair two individuals in this population to generate a single child genome, until a certain number of children have been generated. We then move forward one generation and repeat this process. After each generation, we measure the linkage disequilibrium and compare it to the linkage disequilibirum in a human population used to generate the linkage maps. In the initial population consisting of completely random genomes there is no linkage disequilibrium that resembles a real population; our simulation algorithm then introduces linkage due to random but non-uniformly distributed recombination events so that after several generations the linkage disequilibrium should approximate the linkage found in a human population.

Figure 4 shows the correlation value for the first seven generations. The correlation increases as we move from one generation to the next, and a strong correlation with LD in a human population emerges after only a few generations.

**Figure 4:**
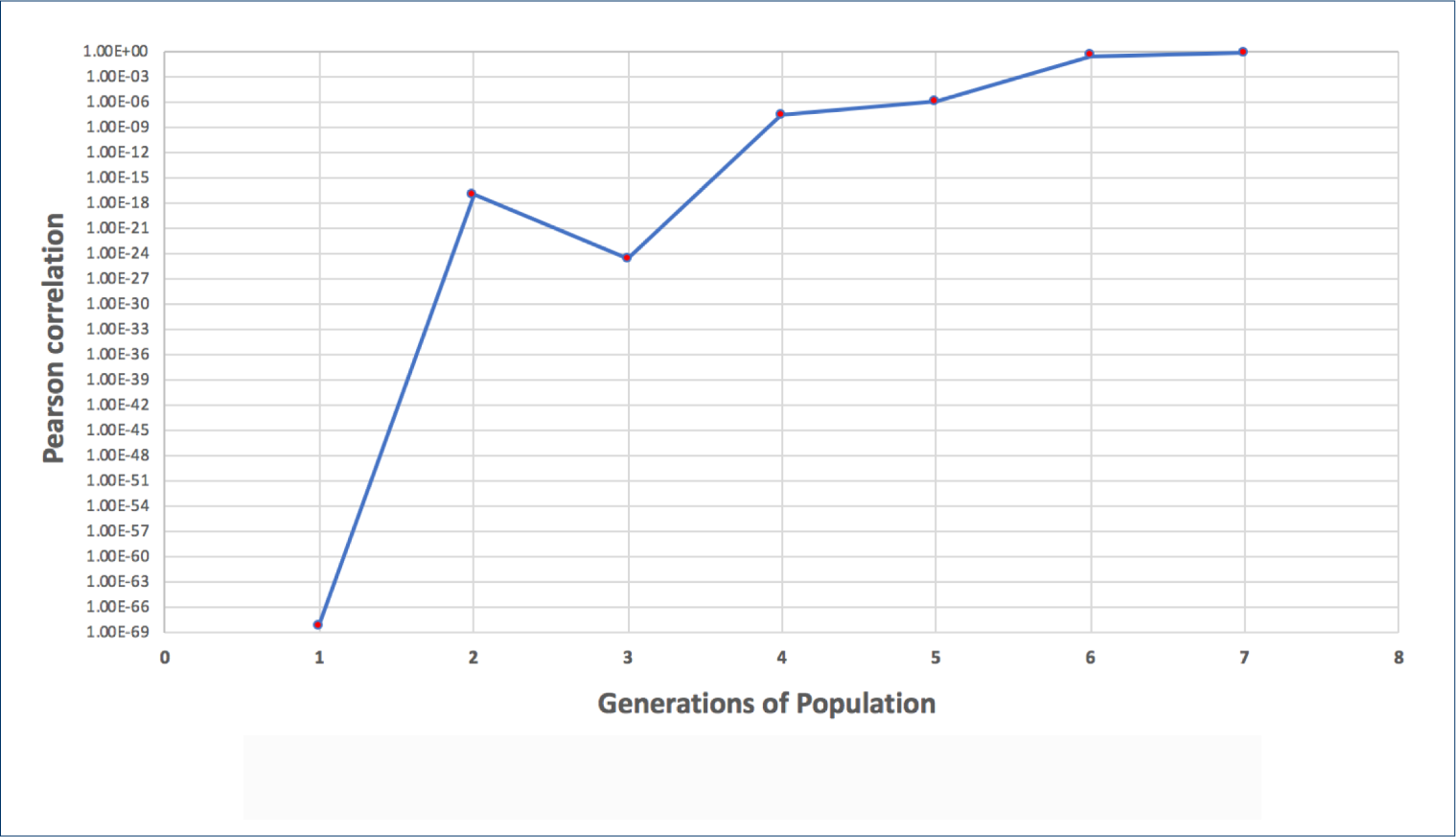
Linkage Disequilibrium Evaluation Result

## Implementation

### Databases for variant annotation

We use four databases (see Table 1) for annotation of genomic variants:

- ClinVar [18] is a database of genomic variants and the interpretation of their relevance to diseases. It identifies the relationships among medically important variants and phenotypes. The variations contained in this database are in VCF format, and ClinVar contains a mixture of variations asserted to be pathogenic as well as those known to be non-pathogenic, with regards to their clinical significance. However, our work focused on the pathogenic and likely pathogenic variants. Therefore, as a result of this restriction, we obtained 84,536 variants out of 396,647 SNPs.
- GWAS [19] is a statistical method that determines the associations between SNPs and particular traits or disorders. The GWAS Catalog [26] now contains over 2,500 unique SNP–trait associations, i.e., associations between single nucleotide variants and phenotypes or diseases. In the GWAS Catalog, we find information about variants (in particular their genomic position) and an association with a (usually) complex diseases (a complex disease is a disease that is multi-factorial and may, for example, be associated with many variants each of which modify disease risk). We use 69,460 variants from the GWAS Catalog in our work.
- DIDA [20] is a database that provides comprehensive information on the genes and associated genetic variants which are associated with digenic diseases (a disease follows digenic inheritance if particular genotypes in exactly two genes explain the disease or phenotype in a patient [27]). DIDA includes 213 digenic combinations which are composed of 364 distinct variants, that involved in 44 digenic diseases [20].
- PharmGKB [21] investigates the association of genetic variation and drugs efficiency. PharmGKB contains pharmacogenetic information related to 3,070 variants. From PharmGKB, variants can be associated with different drug responses.

**Table 1:** Used Databases

| Database | Purpose | Number of Variants |
| --- | --- | --- |
| ClinVar | Mendelian Diseases | 84,536 |
| GWAS Catalog | Complex Diseases | 69,460 |
| PharmGKB | Drug response | 3,070 |
| DIDA | Digenic diseases | 364 |

We used the four databases for annotation (dated 30 Aug, 2018), using the reference genome GRCh37 as the main genomic variants set. However, since some of the databases(ClinVar, GWAS Catalog, and DIDA) update regularly, we identified variants from these databases remotely for the annotation.

For identifying the candidate disease variants with the ClinVar database, we downloaded mode of inheritance (MOI) for diseases from the Human Phenotype Ontology (HPO) database []. As a result, we obtained a total of 6,843 MOI records that we classified into ‘Recessive’, ‘Dominant’, and ‘Others’. We then use the genotype (homo- or heterozygote) of a variant in a VCF files to annotate an individual as affected by a disease or carrying a (heterozygote) disease variant.

### Pathogenicity Prediction

A pathogenic variant is a genetic variant that enhances an individual’s likelihood to develop a particular disorder. The development of the disease symptoms is more likely to appear in the individual when such a variant (or mutation) is inherited. MCAP [22] is a pathogenicity predictor for the rare missense variants within the human genome. It is tuned to the high sensitivity that is required in a clinical context and combines pathogenicity scores of several other tools (including Polyphen-2 [28], SIFT [29] and CADD [30]) within a novel machine learning models and additional features. We use MCAP to predict pathogenicity in all the variants in a VCF file. We use the ANNOVAR tool [31] to perform the annotations.

### Annotating Variants

The annotation algorithm for our tool VSIM uses VCF files directly. The VCF file must include at least the following fields for each variant: chromosome number (#CHROM), position (POS), reference alleles (REF) and alternate alleles (ALT), and information (INFO). We annotate the variants with information related to the databases which we show in Table 1.

### Simulation

We implemented the simulation based on the Real Time Genomics (RTG) simulation tool [32]. RTG provides a blueprint platform for genomic analysis. RTG tools is delivered as an executable with multiple commands executed through a command line interface. RTG, among others, supports the generation of child genomes from two VCF files that represent parents, and contains parameters that allow the specification of the number of recombinations per chromosome as well as the addition of random novel mutations in children. However, RTG’s simulation algorithm for child genomes only supports a completely random recombination of VCF files; we modified the source code accordingly.

We used a set of precomputed recombination rate maps for human genome build 37 [25] to determine recombination probabilities. We converted the recombination rate (with recombination rate measured in cM) to recombination probability using formula 1, following [33]. Formula 1 provides us with a recombination probability for each chromosomal position in the human genome. We use this probability distribution to draw *n* or *m* times (*n* and *m* are parameters that determine the number of cross-overs per chromosome in males and females, respectively) from each chromosome to decide the location of a crossover.

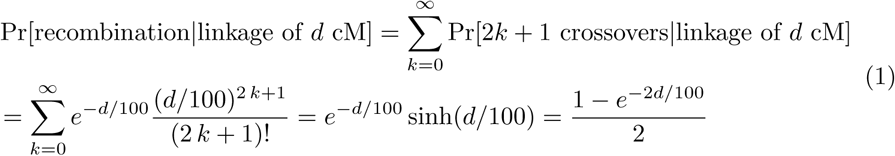

In order to run, the simulator requires two VCF files as an input (representing the mother and father genotype information). Then, the algorithm combines them into a single VCF file. After the combination of VCF files, we generated a population of simulated children (the default number is 100) using our simulation algorithm that accounts for the recombination probabilities. For premarital testing, we can then repeat the step of annotating variants in the resulting simulated children, as before, and generate summary statistics of how many individuals in the simulated cohort are carrying certain disease-associated variants. This summary statistics (and individuals within the simulated cohort) can then be visualized similarly to our visualization of individual VCF files. Algorithm 1 illustrates the procedure that we followed for the simulation.

### Visualization

Genomes and chromosomes are often represented visually through the use of “ideograms”, i.e., a schematic representation of chromosomes, and it is used to show the relative size of the chromosomes and their characteristic patterns. We use the `Ideogram.js` [34] Javascript library for visualizing chromosomes, and we overlay the visual representation of each chromosome with the information obtained from annotating variants in a VCF file.

After obtaining the information related to all the chromosomes’ positions, the next step is to parse genomic features (chromosome, annotation, start and stop of a coding region) from a General Feature Format 3 (GFF3) file which contains the NCBI human genome version 37. We represent the resulting information of variants annotation in JSON, and this file represents the final output of the visualization viewer data Additionally, it contains all the data that is used by the client side in `Ideogram.js`.

#### Algorithm 1: Simulations Algorithm

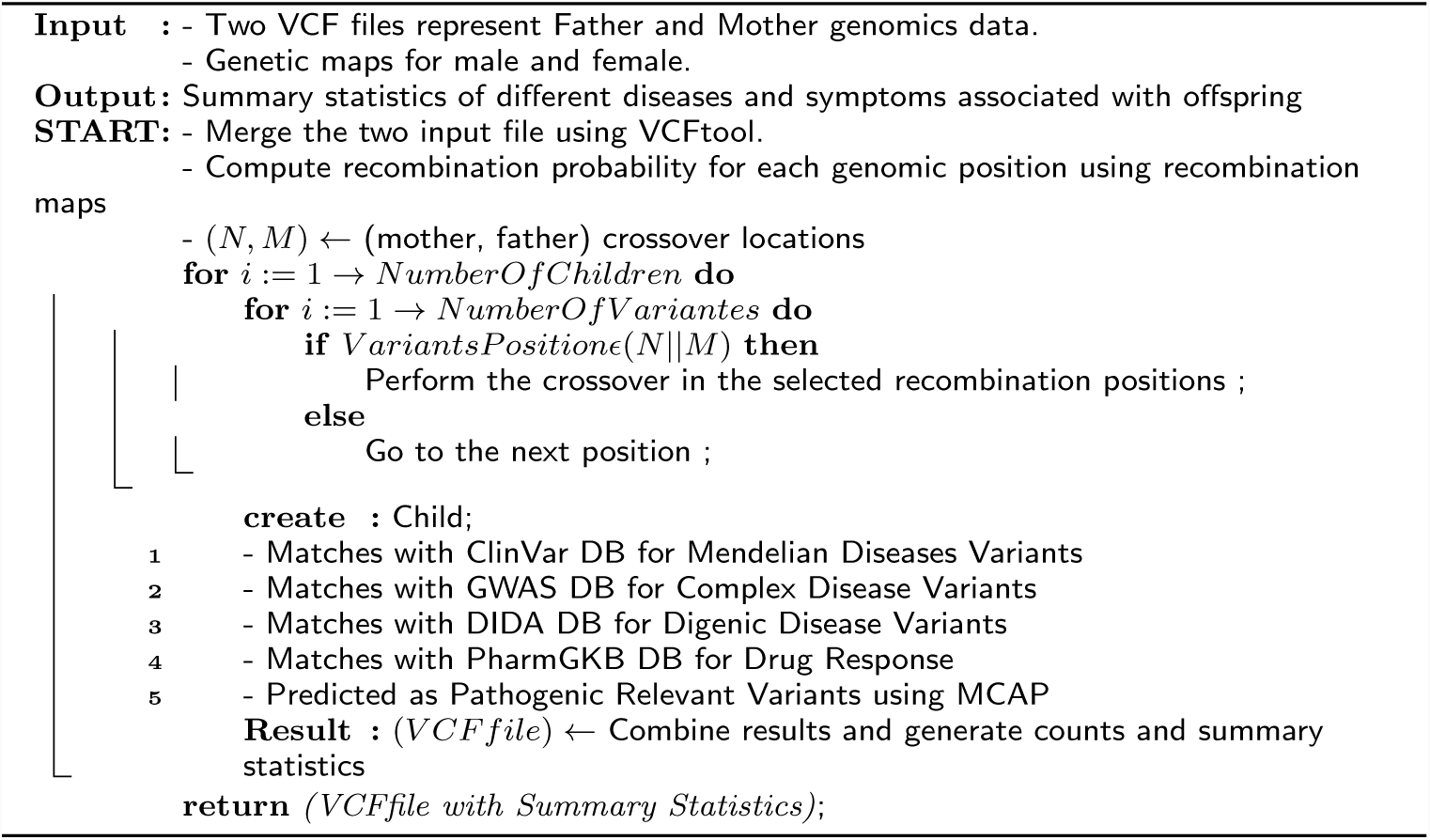

Figure 5 provides an overview of our overall workflow and Figure 6 summarizes the algorithm for annotating and visualizing individual VCF files.

**Figure 5:**
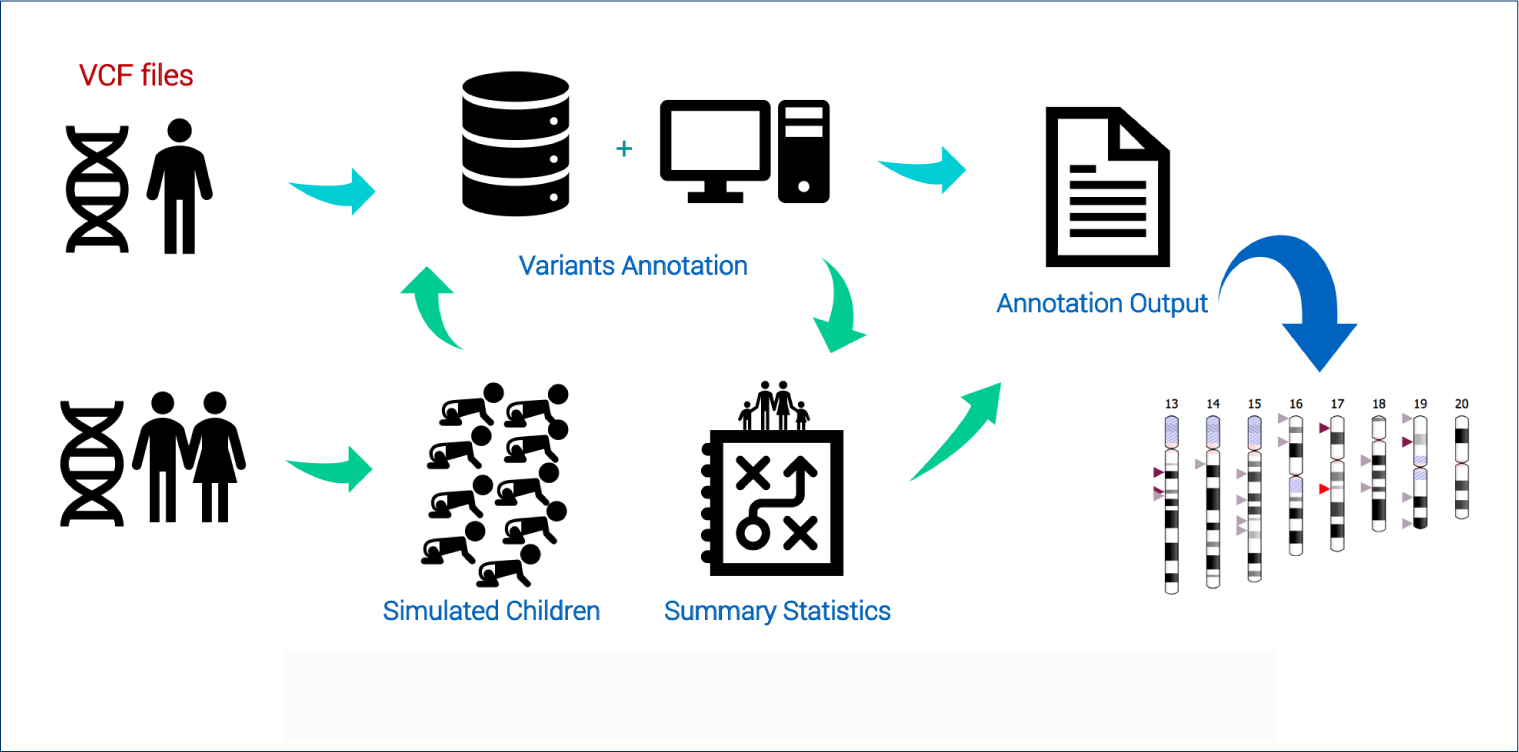
High-level overview over VSIM work-flow

**Figure 6:**
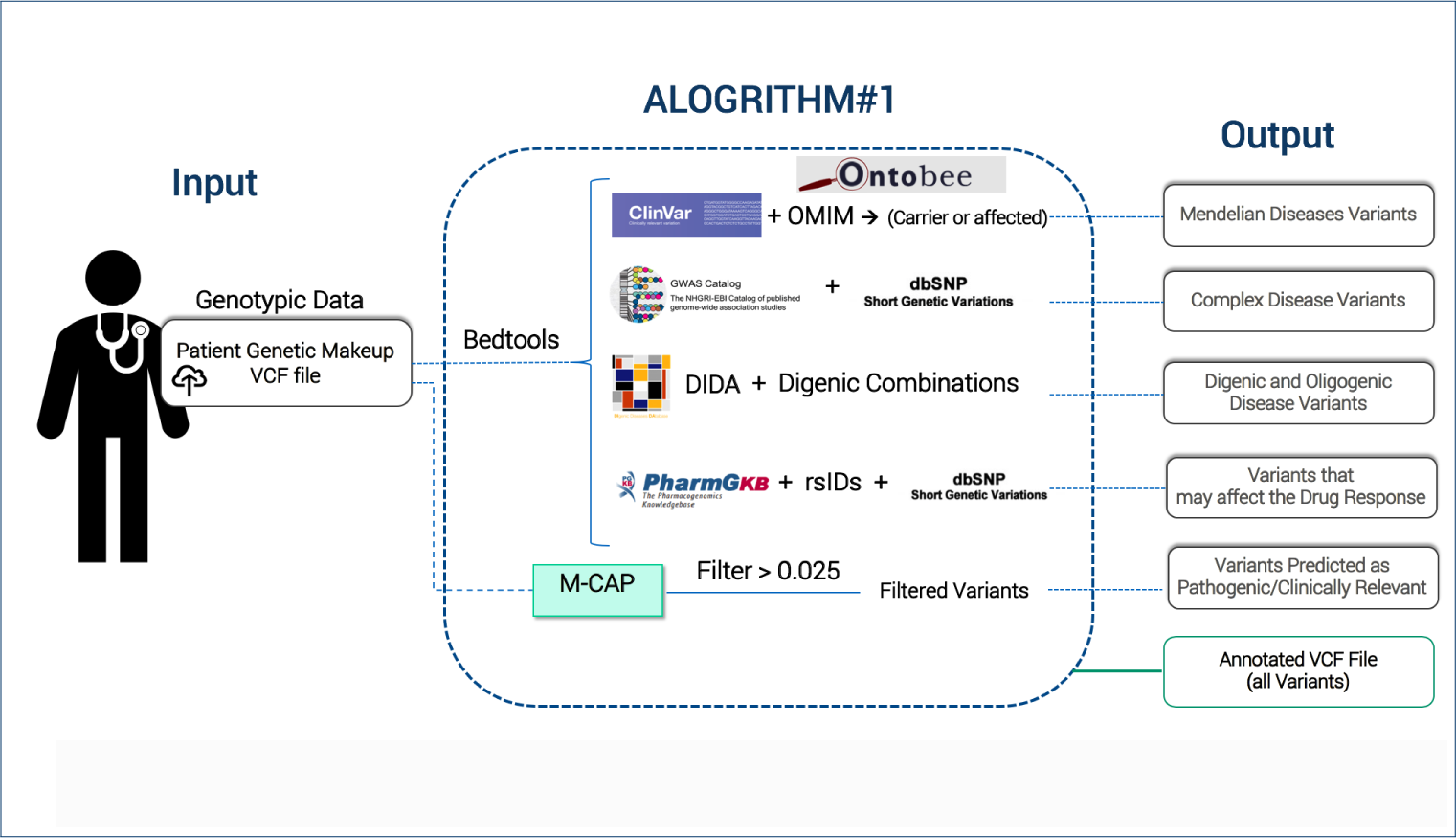
The workflow for analyzing genomic sequence data of individuals

## Discussion

We developed VSIM as a web server that aims to provide an interpretation of individual genomes. Underlying VSIM is a novel genome simulation algorithm that accounts for non-uniformly distributed recombination rates and can be used to create linkage disequilibrium in simulated populations. VSIM can use this simulator to help predict, and to provide a general overview of the potential diseases that might be associated with children.

While this approach is applicable to any disease, its main utility only arises with diseases that are associated with more than one genomic locus. For pre-marital testing, VSIM has several limitations, including the limited number of databases for annotation of genomic variants, its lack of consideration for X- or Y-linked phenotypes, and limited number of polygenic sites and risk scores (mainly coming from known GWAS studies). In the future, VSIM can be extended with additional information about effect sizes of variants and combinations of variants in particular for oligogenic and polygenic disease.

However, the simulator underlying VSIM could also be used as a tool for the study of genetic associations of diseases as well as correlation between different disease-associated loci and their progression within a population. Its application may therefore go beyond premarital testing or interpretation of genomics.

## Conclusions

We have developed VSIM, a server that can be applied in clinical environments for visual interpretation of whole exome or whole genome sequences of individuals. VSIM visualizes genomic features associated with Mendelian, digenic, and complex diseases as well as loci associated with particular pharmacogenomics. Using an accurate genomic simulator, VSIM can also estimate the likelihood that children of two individuals will develop a certain phenotype. VSIM is freely available both as source code and using a Docker container.

## Availability and requirements

- Project name: VSIM
- Project home page: https://github.com/bio-ontology-research-group/VSIM
- Programming language: Python, Javascript, Java
- Operating system(s): Platform-independent
- License: 2-clause BSD license

